# Control of archetype BK polyomavirus miRNA expression

**DOI:** 10.1101/2020.10.20.347229

**Authors:** Wei Zou, Gau Shoua Vue, Benedetta Assetta, Heather Manza, Walter J. Atwood, Michael J. Imperiale

**Affiliations:** Department of Microbiology and Immunology, University of Michigan, Ann Arbor, MI 48109; Department of Molecular Biology, Cell Biology, and Biochemistry, Brown University, Providence, RI 02912

## Abstract

BK polyomavirus (BKPyV) is a ubiquitous human pathogen, with over 80% of adults worldwide persistently infected. BKPyV infection is usually asymptomatic in healthy people; however, it causes polyomavirus-associated nephropathy in renal transplant patients and hemorrhagic cystitis in bone marrow transplant patients. BKPyV has a circular, double-stranded DNA genome that is divided genetically into three parts: an early region, a late region, and a non-coding control region (NCCR). The NCCR contains the viral DNA replication origin and cis-acting elements regulating viral early and late gene expression. It was previously shown that a BKPyV miRNA expressed from the late strand regulates viral large T antigen expression and limits the replication capacity of archetype BKPyV. A major unanswered question in the field is how expression of the viral miRNA is regulated. Typically, miRNA is expressed from introns in cellular genes but there is no intron readily apparent in the BKPyV from which the miRNA could derive. Here we provide evidence for primary RNA transcripts that circle the genome more than once and include the NCCR. We identified splice junctions resulting from splicing of primary transcripts circling the genome more than once, and Sanger sequencing of RT-PCR products indicates that there are viral transcripts that circle the genome up to four times. Our data suggest that the miRNA is expressed from the intron of these greater-than-genome size primary transcripts.

## Introduction

BK polyomavirus (BKPyV), a member of the polyomavirus family, is a ubiquitous human pathogen with over 80% of adults worldwide persistently infected (1, 2). Infection with BKPyV is generally thought to be asymptomatic in healthy individuals despite periodic episodes of reactivation and viral shedding in the urine. In renal transplant patients, reactivation of BKPyV in the donor-derived kidney causes polyomavirus-associated nephropathy (PVAN), and in the bladder of bone marrow transplant patients causes hemorrhagic cystitis (3–5). There is increasing evidence that BKPyV plays a role in the development of urothelial and bladder cancer in transplant recipients who have had BKPyV viremia or PVAN (6–11). There are currently no FDA-approved drugs to treat BKPyV-associated diseases. The primary treatment for interstitial cystitis is palliative, and for PVAN is lowering the dosage of immunosuppressants to allow the immune system to clear the virus, which inevitably increases the risk of acute rejection and graft failure (12).

BKPyV has a double-stranded circular DNA genome which is divided genetically into the early region, late region, and the noncoding control region (NCCR). The early region encodes the large T and small T antigens (TAg and tAg), which are important for BKPyV replication, and a truncated TAg that induces expression of cellular APOBEC3B (13, 14). The late region is expressed from the complementary strand of the genome and encodes the viral capsid proteins VP1, VP2, and VP3, and an auxiliary protein called agnoprotein. Agnoprotein has been reported to facilitate BKPyV replication by disrupting the mitochondrial network and mitochondrial membrane potential, allowing evasion of innate immunity (15), as well as playing a role as a viroporin in viral egress (16, 17). The NCCR contains the cis-acting elements that are required for regulating DNA replication, including the origin of replication, and transcription of the early and late regions. Based on the NCCR structure, there are two forms of the BKPyV genome, archetype virus and rearranged variants. The NCCR structure of archetype virus is arbitrarily divided into five sequence blocks termed O, P, Q, R, and S, while the NCCRs of rearranged variants are generally characterized by duplications and/or deletions of the P, Q, R, and S blocks (18, 19). The O block is essential and cannot be deleted because it contains the origin of DNA replication. The archetype virus is thought to be the transmissible form of the virus because it is found in both healthy people and in diseased patients, while rearranged variants are usually isolated from patients with BKPyV disease (20, 21).

BKPyV also encodes a miRNA on the late gene strand that targets early mRNAs for degradation, as is the case for other polyomavirus miRNAs (22–26). Our previous study showed that the BKPyV miRNA plays an important role in limiting archetype virus replication but not rearranged variant replication: a miRNA mutant archetype virus showed a much higher replication capability than the wild type virus. We proposed a model in which the miRNA’s function is mediated through the balance of regulatory elements in the NCCR that control relative expression of the miRNA and the early genes (23).

Although it is clear that the BKPyV miRNA plays an important role in controlling early gene expression, how the expression of the miRNA is regulated is not known. Previous studies have shown that most cellular miRNAs are expressed from introns of protein coding genes. However, the BKPyV miRNA is not encoded in an obvious intron. In the present study, we provide evidence that the miRNA is expressed from a putative intron that is generated by transcription that circles the viral genome more than once.

## Materials and Methods

### Cell culture

Primary human Renal Proximal Tubule Epithelial (RPTE) cells (Lonza) were maintained and passaged in REGM medium (REGM/REBM, Lonza) as described previously. RPTE cells obtained from ATCC (American Type Culture Collection) were cultured in Renal Epithelial Cell Basal Medium (RECBM) supplemented with Renal Epithelial Cell Growth Kit. 293 cells and 293TT cells (14) were cultured in Dulbecco’s modified Eagle medium (DMEM) with 10% FBS and 100 units/mL penicillin, 100 μg/mL streptomycin. All cells were cultured in a humidified incubator at 37°C with 5% CO2.

### Virus infection

Archetype and rearranged BKPyV (Dik and Dunlop, respectively) were purified and titrated as described previously (27). RPTE cells at passage 5 were plated in 12-well plates for infection. Cells were prechilled for 15 min at 4°C and infected with Dik or Dunlop at a MOI of 2 fluorescence-forming unit (FFU)/cell. A 400 μl volume of the diluted virus was added to each well and incubated at 4°C for 1 h with gentle shaking every 15 min. The virus was aspirated off and the cells were incubated with fresh REGM medium.

### Transfection

For the experiments comparing circular and linearized plasmids, pGEM-Dik-LT/O, that contains the NCCR, late coding region, and miRNA coding region, and the GFP expression vector pEGFP-N1 were linearized with DraIII and treated with alkaline phosphatase before purifying with a PCR purification kit (Qiagen). Then 500 ng each of circular or linearized pGEM-Dik-LT/O and pEGFP-N1 were co-transfected into 293 cells with Lipofectamine 2000 reagent according to the manufacturer’s instruction and the medium was replaced with fresh DMEM four hours after transfection. Two days post-transfection, total RNA was extracted using the Direct-zol RNA MiniPrep kit (ZYMO Research, USA). GFP expression was visualized using an Olympus microscope, and the number of GFP-positive cells were determined using ImageJ (NIH).

### RT-PCR amplification of the NCCR

RPTE cells were infected with Dik and Dunlop as described above. Total RNA was extracted using the Direct-zol RNA MiniPrep kit (ZYMO Research, USA) and treated with DNaseI (Promega). Reverse transcription (RT) was performed using an NCCR late strand-specific primer NCCR-r (5’-GGTACCGGCCTTTGTCCAGTTTAACT-3’) (Figure 1A), followed by PCR to amplify the NCCR using primers NCCR-f (5’-CTCGAGTTGCAAAAATTGCAAAAGAATAGG-3’) and NCCR-r. pGEM-Dunlop plasmid, which contains the entire Dunlop genome, and water were used as positive and negative controls, respectively. The PCR products were electrophoresed in a 2% agarose gel and visualized using a Bio-Rad gel system (Bio-Rad).

**Figure 1.**
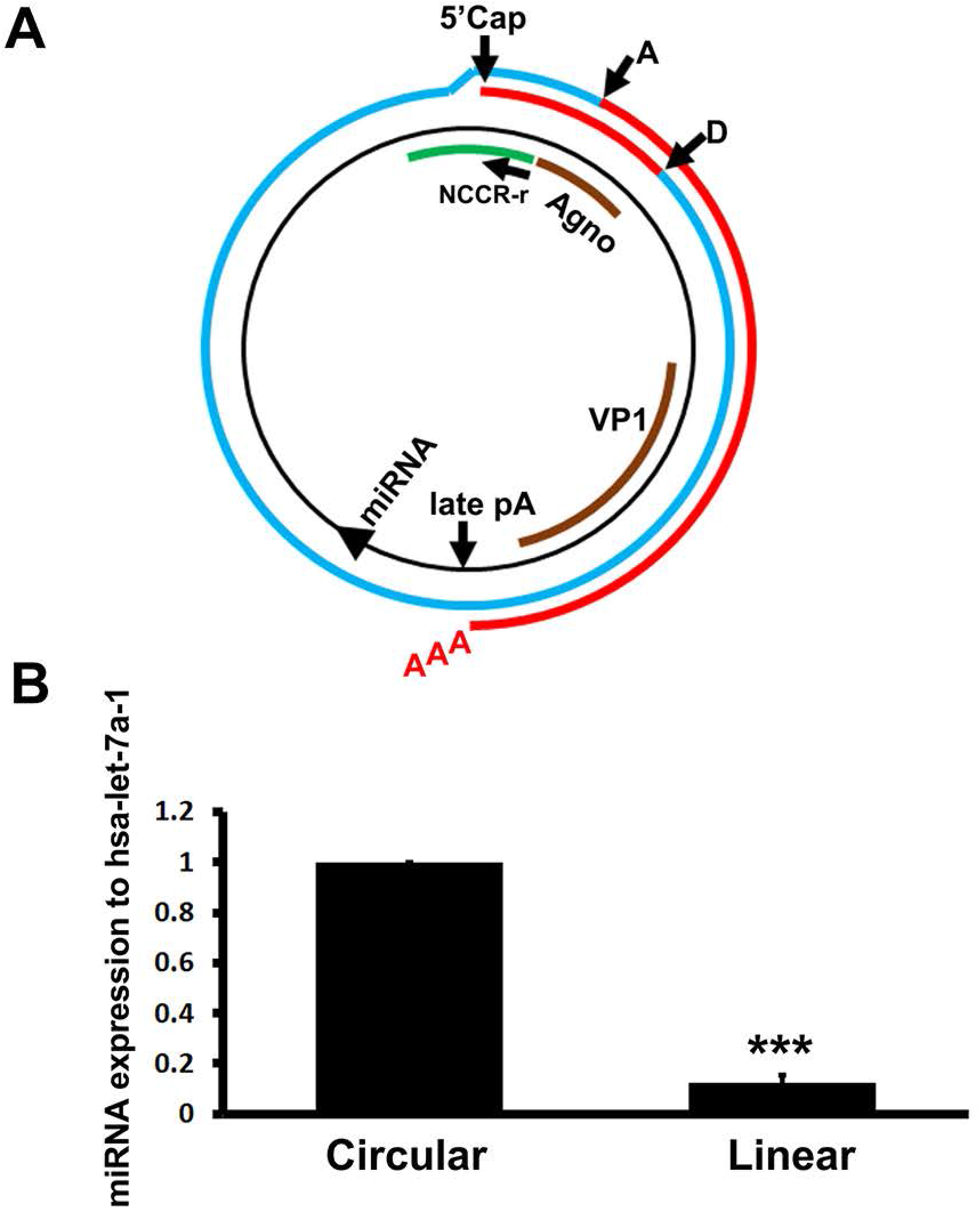
A circular viral template expresses higher levels of miRNA than a linear template. **(A)** Model of miRNA biogenesis, showing the pre-miRNA coming from the intron (blue line) of a primary transcript that is longer than the genome. The exon in this example (red), when spliced, forms a VP1 mRNA. D, splice donor; A, splice acceptor; Late pA, late polyadenylation signal site. The mature miRNA is processed from the putative intron. The location of the primer NCCR-r, which was used to perform reverse transcription to confirm transcription of the NCCR, is shown **(B)**. Total RNA was extracted from transfected 293 cells and assayed for BKPyV miRNA expression using stem-loop RT qPCR. The results are normalized to the cellular miRNA control, hsa-let-7a-1. The circular template value was arbitrarily set to 1. *** p<0.005.

### Quantitative stem-loop qPCR

miRNA stem-loop qPCR was performed and the miRNA expression level was normalized to human miRNA hsa-let-7a-1 using the 2^−ΔΔC(t)^ method as previously described (23, 28).

### Next generation sequencing (RNA-seq) analysis

RPTE cells were infected with BKPyV Dunlop at an MOI of 0.03 FFU/cell as follows. Virus was diluted in culture media and cells were infected for 2 hours. The inoculum was aspirated and cells were incubated with fresh medium. Cells were fed every 2 days and medium replaced at 6 days. Total RNA was harvested from infected and mock-infected cells at 3 days post infection (dpi), 6 dpi, and 9 dpi with a RNeasy Mini Kit (Qiagen) with DNase treatment. Quantification was performed using a Nanodrop 2000c and the quality of the RNA was assessed with an Agilent 2100 Bioanalyzer. Libraries were prepared with a TruSeq Stranded Total RNA Sample with Ribo-Zero Prep Kit and the RNA sequencing read type was 2×50bp. Library preparation and RNA sequencing were performed by Beckman Coulter Genomics (Genewiz). The number of Quality Control (QC)-passed reads varied between ~60 and 89 million across all samples (29). Hisat2 2.1.0 was used to map RNA-seq reads against the Dunlop full-length genome using default parameters. The BKPyV reads were extracted using an in-house-developed script. The alignment data were used to compute coverage by igvtools count (https://software.broadinstitute.org/software/igv/igvtools). Transcript abundance estimation of NCCR, early genes, and late genes was conducted with StringTie 1.3.5 software and is reported in Fragments Per Kilobase of exon per Million fragments mapped (FPKM).

### TA cloning and Sanger sequencing

Specific primer pairs for confirming the most abundant splice junctions identified from the RNA-seq data were used to amplify cDNA transcribed from RNA isolated from Dik-infected RPTE cells using SuperScript™ III Reverse Transcriptase kit (Invitrogen). The sequences of the primers are primer 1 (P1) 5’-GACTCTGTAAAAGACTCCTAG-3’ and primer 2 (P2) 5’-CTAAAATAAAAATAAAAATCCTCTG-3’ (Figure 3A). The amplified band was purified using a Qiagen gel purification kit and TA cloning was performed using a TOPO™ TA cloning™ kit (Invitrogen). 12 clones were randomly picked for Sanger sequence analysis by the Advanced Genomics Core at the University of Michigan.

## Results

### A circular template expresses higher levels of the miRNA than a linear template

It is known that miRNAs are usually expressed from introns of protein-coding genes (30). The BKPyV miRNA is not encoded in an obvious intron since it maps downstream of the late polyadenylation site (Figure 1A). However, it is possible that the miRNA could be expressed from an intron of a primary RNA transcript that results from transcription initiating in the NCCR and circling the genome more than once (Figure 1A). Such greater-than-genome size primary transcripts are known to be produced from the related murine polyomavirus (MuPyV) (31, 32). If this is the case, the prediction is that linearizing the genome would eliminate such transcripts and reduce miRNA expression. To test this hypothesis, we transfected 293 cells with pGEM-Dik-LT/O, which is a genomic clone of the BKPyV NCCR, late region, and sequences downstream of the late poly(A) site that encode the miRNA; or the same plasmid linearized by DraIII digestion (whose recognition site is located in the vector backbone and does not cut within the viral sequences), to mimic circular and linear viral genomes, respectively. A circular or linearized GFP-expressing plasmid was co-transfected with the circular and linear viral plasmids, respectively, to monitor transfection efficiency. The results showed that miRNA expression is significantly lower from the linear viral plasmid than from the circular plasmid (Figure 1B).

### The NCCR region is transcribed

To demonstrate that RNA polymerase circles the viral genome, we infected RPTE cells with archetype BKPyV and extracted RNA 3 days post infection. Reverse transcription was performed using a primer that is complementary to the late strand of the NCCR (Figure 1A), followed by PCR to amplify the NCCR. We detected a specific band in archetype-infected RPTE cells but not mock-infected cells or when the RT step was omitted, indicating that late strand viral RNA transcripts containing the NCCR are produced (Figure 2A). Viral RNA containing the NCCR was also observed in rearranged BKPyV-infected RPTE cells (Figure 2B). To confirm that the NCCR was transcribed, we analyzed RNA-seq transcriptome data from rearranged virus-infected RPTE cells. The number of reads mapping to the viral genome increased over time, from approximately 5% (3.5 million reads) at 3 dpi to almost 30% (26 million) of the total reads at 9 dpi. A significant number of reads spanning the NCCR were detected (Figure 2C, Table 1). The overall NCCR late strand steady state expression level is approximately 10% of that of the late viral coding sequences (Table 2).

**Figure 2.**
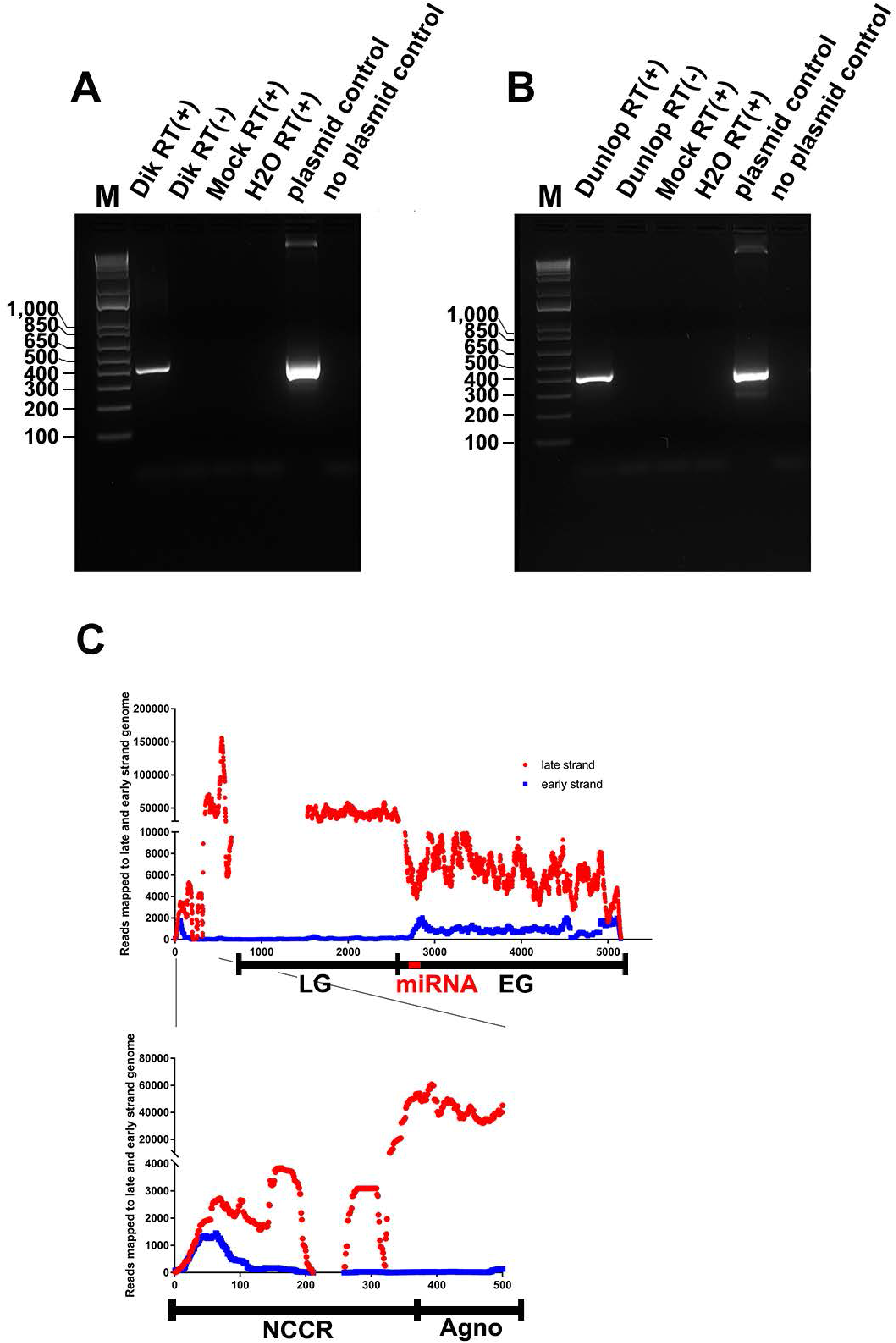
Viral RNA transcripts contain the NCCR region. RPTE cells were infected with archetype Dik virus **(A)** or rearranged variant Dunlop virus **(B)**, or mock infected. Total RNA was extracted 3 dpi. Reverse transcription (RT) and PCR amplification with specific NCCR primers (Methods and Figure 1A) were performed and the PCR products were electrophoresed on a 2% DNA agarose gel. Representative gels from three independent repeats with each virus are shown. **(C).** RNA-seq read coverage of the Dunlop genome. One representative sample was shown. Read coverage on the late strand (red) and early strand (blue) are shown graphed along the length of the viral genome. The locations of the non-coding control region (NCCR), agnoprotein gene (Agno), early genes (EG), late genes (LG), and miRNA gene are indicated. The enlarged area showed the coverage for NCCR and Agno regions.

**Figure 3.**
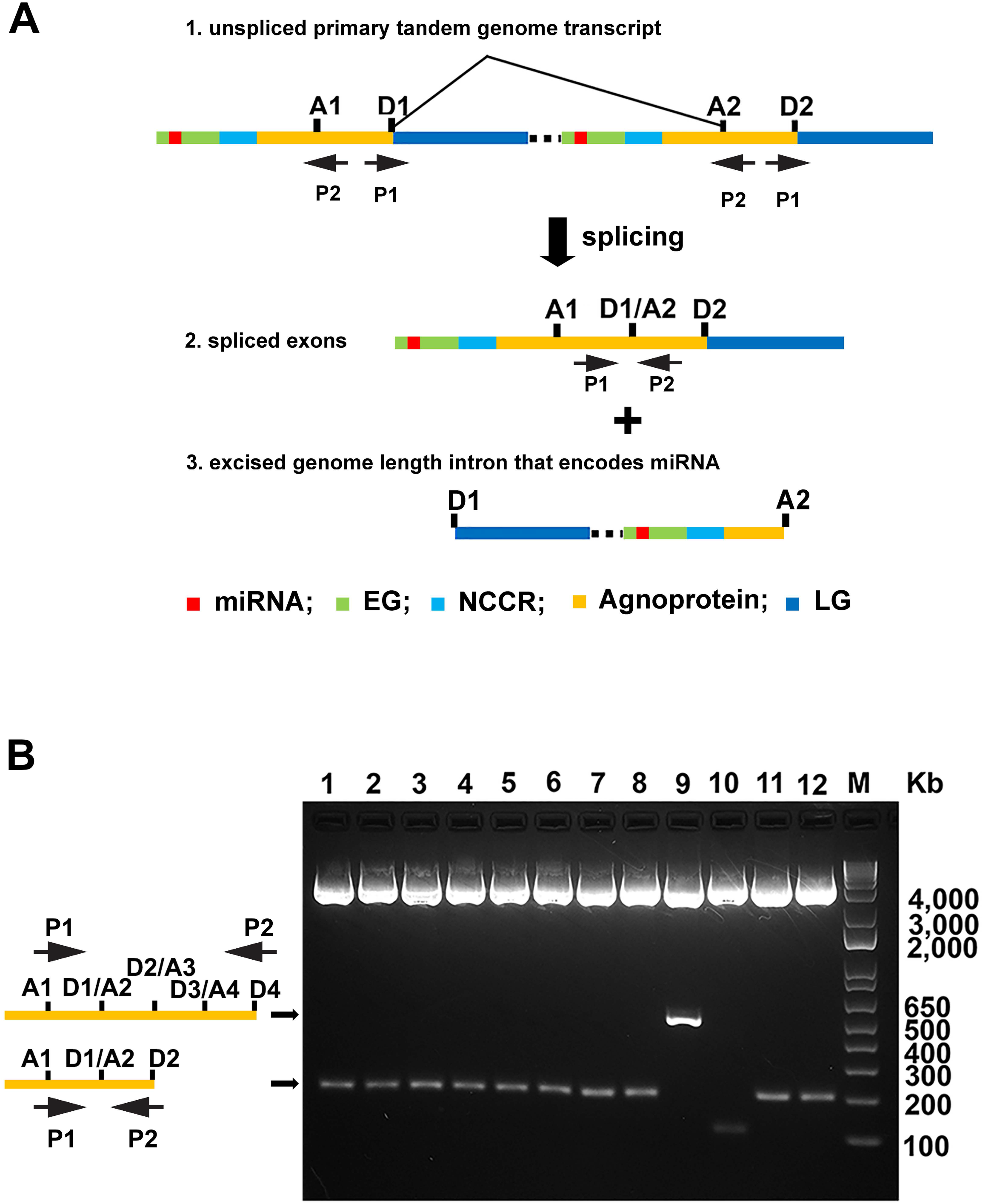
Evidence for viral primary mRNAs that circle the genome more than once. **(A).** (1) linear representation of a primary tandem genomic transcript resulting from the RNA polymerase transcribing the genome more than once. The relevant genomic regions are shown in different colors (not to scale), and the locations of putative splice donors (D) and acceptors (A) based on the RNA-seq data are shown. A1 and D1 are in the first tandem copy, and A2 and D2 are in the second copy. Also shown are the two PCR primers (1 and 2) used to analyze the RNA products. cDNA from this unspliced RNA would not be amplified by primers 1 and 2 in our assay because the product would be > 5 Kb. (2) Spliced product resulting from excision of a genome-sized intron. Primers P1 and P2 can now amplify cDNA from this RNA, yielding a product of 99 bp, or larger in size if the transcript circled the genome more than twice and was subsequently spliced. 3) Intron resulting from splicing to give the product in (2). **(B).** TA cloning to confirm the splice junctions detected in the RNA-seq data. Specific primers shown in (A) were used to amplify the cDNA transcribed from RNA of archetype-infected RPTE cells. PCR products were cloned into pCR2.1-TOPO vector, and 12 random clones were chosen for DNA minipreps. The DNA was digested with BamHI and XbaI and electrophoresed on a 2% DNA agarose gel. The sizes of the bands are 94 bp longer than the cDNA insert due to the location of the restriction sites used to excise the inserts. The diagrams to the left of the gel indicate the structure of the spliced products represented by the different sized bands. Lane 10 is empty pCR2.1-TOPO vector digested with BamHI and XbaI, which produces a 94 bp band because there is no insert.

**Table 1.**
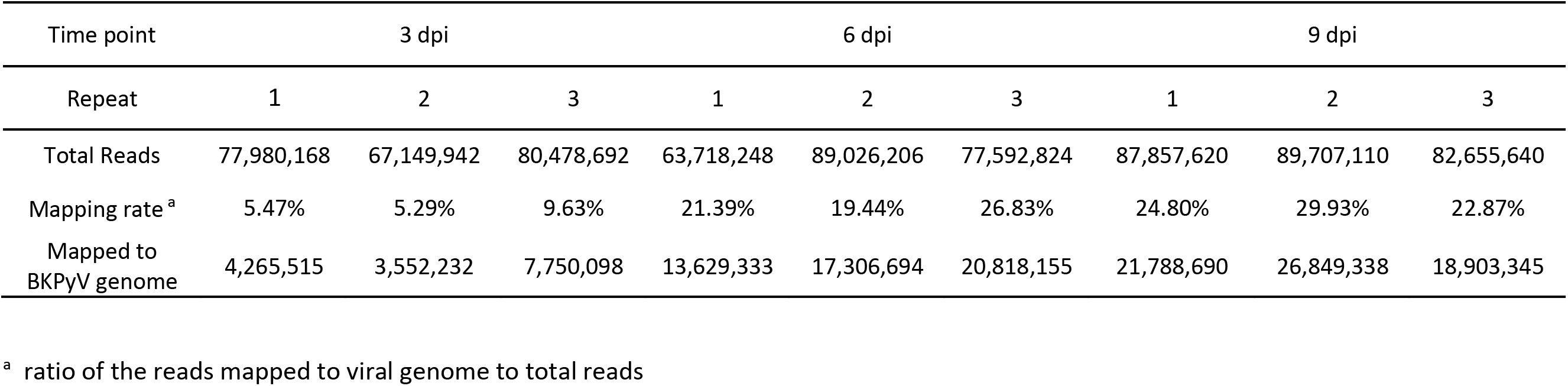
Summary of reads mapped to viral genome from RNA-seq data

**Table 2.** NCCR expression levels

| Time point | Experiment | Late region<br>(RPKM) | NCCR<br>(RPKM) | (NCCR/late)% | mean ± SD<br>(NCCR/late)% |
| --- | --- | --- | --- | --- | --- |
| 3 dpi | 1 | 17,304 | 1,986 | 11.48 | 11.48 ± 0.67 |
|  | 2 | 17,033 | 2,095 | 12.30 |  |
|  | 3 | 31,035 | 3,305 | 10.65 |  |
| 6 dpi | 1 | 66,143 | 6,561 | 9.92 | 9.47 ± 0.39 |
|  | 2 | 60,987 | 5,815 | 9.53 |  |
|  | 3 | 84,661 | 7,582 | 8.96 |  |
| 9 dpi | 1 | 75,326 | 7,055 | 9.37 | 9.16 ± 0.88 |
|  | 2 | 100,071 | 7,992 | 7.99 |  |
|  | 3 | 73,899 | 7,480 | 10.12 |  |

### Identification of viral transcripts resulting from RNA polymerase transcribing the genome more than once

The detection of viral transcripts containing the NCCR by both RT-PCR and RNA-seq supported the possibility of production of a primary RNA transcript resulting from transcription circling the genome more than once (Figure 1A). Such a primary transcript could be spliced in a manner similar to what has been reported for MuPyV, in which a genome-sized intron is removed. To test this, we further analyzed the RNA-seq data of Dunlop-infected RPTE cells for the presence of splice junctions that could only be produced by splicing as indicated in Figure 1A. To facilitate this analysis, we aligned the reads to a template consisting of two tandem copies of the viral genome. We identified many reads containing splice junctions that support the existence of primary transcripts which circle the genome more than once (Table 3). To produce such junctions, an intron containing the miRNA coding sequence would have had to have been excised. The most abundant splice junctions are ones in which the donor site is in the first copy of the genome while the acceptor site is located in the second copy of the genome, which only could occur if the primary transcripts circled the genome more than once and splicing occurred from a donor site such as D1 to an acceptor site such as A2 (Figure 3A, Table 3). Overall, the splice junctions arising from transcripts circling the genome more than once occur at different sites on the viral genome and are enriched within the agnoprotein and VP1 genes. We also observed an increase in the number of transcripts containing these splice junctions over time, although their relative abundance compared to the authentic VP1 splice junction (D3041/A3974) remained constant (Table 3).

**Table 3.** Summary of splice junctions of viral RNA

|  |  | 3 dpi <sup>c</sup> |  |  | 6 dpi |  |  | 9 dpi |  |  | location <sup>d</sup> |
| --- | --- | --- | --- | --- | --- | --- | --- | --- | --- | --- | --- |
|  |  | 1 | 2 | 3 | 1 | 2 | 3 | 1 | 2 | 3 |  |
| Donor (D) <sup>a</sup> | Acceptor (A) <sup>b</sup> | BK3DPI1 <sup>c</sup> | BK3DPI2 | BK3DPI3 | BK6DPI1 | BK6DPI2 | BK6DPI3 | BK9DPI1 | BK9DPI2 | BK9DPI3 |  |
| 3,041 | 8,093 (2,940) | 117,947 | 102,833 | 230,080 | 311,641 | 426,286 | 491,182 | 433,764 | 606,023 | 359,356 | Agno |
| 3,014 | 8,117 (2,964) | 20,128 | 18,445 | 40,988 | 61,794 | 80,724 | 95,341 | 82,856 | 149,456 | 93,262 | Agno |
| 3,041 | 8,115 (2,962) | 16,855 | 10,218 | 29,058 | 45,737 | 64,424 | 69,149 | 59,551 | 106,723 | 75,976 | Agno |
| 5,109 | 10,104 (4,951) | 1 | 16,795 | 3 | 70,650 | 96,021 | 168,128 | 164,276 | 205,883 | 2 | VP1 |
| 5,109 | 10,231 (5,078) | 5 | 9 | 37,384 | 1,007 | 55 | 52 | 44 | 135,253 | 97,293 | VP1 |
| 5,000 | 10,114 (4,961) | 2 | 1,945 | 8,738 | 20,236 | 37,697 | 8,229 | 13,706 | 96,622 | 61,331 | VP1 |
| 3,041 <sup>e</sup> | 3,974 | 57,854 | 52,111 | 107,881 | 150,276 | 202,325 | 248,315 | 206,876 | 344,419 | 203,219 |  |

<sup>a</sup> We set the T in the sequence TAAGCCAC, located between the 3' ends of the VP1 and TAg coding sequences of the Dunlop genome, as nucleotide 1, which places the NCCR in the middle of the genome and facilitates comparing the reads to a tandem genome as described in the text (i.e., nucleotide 5154 is the same T in the second copy of the genome).
<sup>b</sup> the location of the acceptor site of the reads containing a splice junction in the second copy of the virus genome. The number in parentheses indicates the location of the equivalent acceptor site in the first copy of the virus genome (note that these acceptor sites in the first copy are located upstream of the donor sites).
<sup>c</sup> The number of the reads for each representative splice pattern is shown.
<sup>d</sup> the coding location of the splice junction in the viral genome. Agno, agnoprotein.
<sup>e</sup> The splice junction D3041/A3974 produces authentic VP1 mRNA.

To confirm the presence of the most abundant splice junction, we designed a PCR primer pair that would specifically amplify RNAs that are spliced from one copy of the tandem primary transcript to the second copy (Figure 3A). We used this pair of primers to amplify cDNA transcribed from RNA of archetype-infected RPTE cells and cloned the PCR products into the pCR2.1-TOPO vector. We identified specific inserts from individual colonies that matched the splice junctions detected in the RNA-seq analysis (Figure 3B, lane 1-8, 11-12). Sanger sequencing of the DNA isolated from individual colonies confirmed the PCR products were perfectly matched with the predicted splicing patterns. Most interestingly, we identified an RT-PCR product whose sequence indicates that the RNA polymerase circled the genome four times (Figure 3B, lane 9).

## Discussion

Most polyomaviruses encode a miRNA from the late strand of their circular genome. Although the genetic location of the miRNA and the target sequence differ among the polyomaviruses, all the miRNAs share in common the ability to regulate expression of the major viral regulatory protein, TAg (23–26, 33). We previously proposed a model we called the promoter balance model to explain how the NCCR regulates viral early gene, late gene, and miRNA expression (23). In archetype virus, the NCCR drives low early promoter activity and high late promoter activity, resulting in a relatively high level of miRNA, which further inhibits viral early gene expression and virus replication. In rearranged variants, the NCCR has weak late promoter activity and high early promoter activity, resulting in a high level of early gene expression and robust virus replication. However, how the viral miRNA was regulated or if there any other promoter sequence regulates miRNA expression is not fully understood. A previous study with Merkel cell polyomavirus (MCPyV) showed that an approximately 100 base pair sequence upstream of the miRNA coding sequence contains a promoter activity that can initiate viral miRNA expression independently of NCCR-initiated transcription (34). Unlike these authors, we did not detect any obvious spike in miRNA reads in our RNA-seq analysis of BKPyV, although our RNA-seq preparations were not optimized for small RNAs. We have also begun to dissect the viral cis-acting sequences that regulate miRNA expression. Our preliminary results suggest that there are sequences both within and outside the NCCR that serve this function (W.Z., unpublished data).

Previous studies have shown that most cellular miRNAs are expressed from intron sequences. However, the BKPyV miRNA is not encoded in an obvious intron. The apparent answer to this dilemma was first demonstrated by the Carmichael lab in their studies of MuPyV. They determined that RNA polymerase II can generate a primary RNA transcript that circles the genome more than once, due to the presence of a weak late strand polyadenylation site (31, 35). Their results showed that genome-length presumptive introns could be spliced out of these primary transcripts that represented tandem copies of the viral genome. Viral late transcripts that circle the genome more than once have also observed more recently in MCPyV-infected PSFK-1 cells (34). Based on these results, we took three approaches to address the question of whether the BKPyV miRNA can be expressed without being present in an apparent intron. First, we showed that a circular DNA template expresses higher levels of the miRNA than a linear template. Second, we detected the NCCR in viral RNA transcripts by both RT-PCR amplification and RNA-seq analysis. Third, we identified splice junctions that could only be generated if primary transcripts circled the viral genome more than once. Interestingly, we found evidence of primary RNAs that circle the genome up to four times, albeit at a low frequency. In the MuPyV study, a sequence they named the leader sequence was found in the mature late gene mRNA (31, 32). This sequence contained multiple tandem copies of an untranslated 57 nucleotide segment located at the 5’ end, which indicated that the primary viral RNA circled the genome more than once. It was also shown that the length of the leader sequence is important for virus viability, which includes regulating both pre-mRNA splicing and stability. In these experiments, precise donor and acceptor sites were used for splicing of the leader sequence and an exact genome length intron was excised. In our study, we found using more sensitive assays that the splice junctions occur at different locations, albeit with enrichment in the agnoprotein and VP1 genes, producing different tandem exons and excised introns. Although back splicing or splicing of circular RNAs also could generate the splice junctions that we detect, we think this is a less likely explanation.

Why the virus uses this complicated way to regulate viral late gene expression is not clear. However, since the coding capacity of small viruses like polyomavirus is limited, the virus could take advantage of this unique method of expressing both proteins and a miRNA despite their genome size constraint. The MuPyV studies showed that the leader-to-leader splicing is also essential for leader-VP1 splicing, which is required for mature VP1 mRNA production (32). Since these splice junctions occur at different locations on the BKPyV primary transcripts, the function of these transcripts is unknown and warrants further study.

Taken together, these results indicate the complexity of the regulation of viral RNA expression. We confirmed that the miRNA is likely expressed from an intron that would be present in viral primary transcripts. The RNA-seq data showed that the expression level of the NCCR is about 10% of the late gene expression level. While this may indicate that only a portion of the RNA transcripts circle the genome more than once, since RNA-seq only measures steady state RNA, the NCCR may be transcribed at a higher rate but the products are not detectable because the putative intron would be degraded after splicing and processing of the miRNA.

Although it is well known that the miRNA from different polyomaviruses share the common ability to regulate expression of viral TAg, which has been proposed to modulate persistent infection (23, 34, 36, 37), the miRNAs may also harbor distinct functions. The SV40 miRNA has been shown to downregulate TAg expression as a means of immune evasion, reducing cytotoxic T lymphocyte susceptibility of infected cells (26). However, a mutant MuPyV virus that does not express the miRNA exhibited the same infection and transformation phenotypes as wild type MuPyV, and the mutant induced an indistinguishable immune response as the wild type virus in mice (24). On the other hand, the MuPyV miRNA seems to control viruria in acutely-infected animals (36). It has also been reported that the BKPyV and JCPyV miRNAs target the cellular stress-induced ligand ULBP3 to allow the virus to escape detection by the immune system (38). In addition, the MCPyV miRNA regulates cellular mRNA expression (39). Illustrating how the expression of polyomavirus miRNAs are regulated will shed light on their function in pathogenesis and oncogenesis.

## Acknowledgements

The authors would like to thank Andrew Tai for critical reading of the manuscript. This work was supported by NIH/NIAID grant AI060584 awarded to M.J.I. and NIH/NINDS grants R01NS043097 and P01NS065719 to W.J.A. Research reported in this publication was also supported by the NIH/NCI under Award Number P30CA046592 by the use of the following Cancer Center Shared Resource: DNA Sequencing Core.

## Notes

### Competing Interest Statement

The authors have declared no competing interest.

